# Benchmarking AlphaSC: A Leap in Single-Cell Data Processing

**DOI:** 10.1101/2023.11.28.569108

**Authors:** Hy Vuong, Tam Luu, Nhat Nguyen, Nghia Tra, Ha Nguyen, Huy Nguyen, Thao Truong, Son Pham

## Abstract

We benchmarked AlphaSC, BioTuring’s GPU-accelerated single-cell analytics package, against other popular tools including Scanpy, Seurat, and RAPIDS. The results demonstrate that AlphaSC operates thousands of times faster than Seurat and Scanpy. Additionally, it surpasses RAPIDS, another GPU-accelerated package from NVIDIA, by an order of magnitude in terms of speed while also consuming considerably less RAM and GPU memory. Importantly, this significant increase in AlphaSC’s performance does not compromise its quality. ^1^

## 1 Introduction

BioTuring has established the world’s largest single-cell database by processing the transcriptomes of hundreds of millions of cells and manually curating and transferring annotations from thousands of single-cell research publications. Processing large single-cell datasets poses a major challenge due to the extensive time and computing resources required. For a dataset with a million cells, standard pipelines demand many hours and high-memory servers. NVIDIA’s RAPIDS package [1], which utilizes GPU, does improve speed but hasn’t reached the level where it enables real-time, interactive exploration and adjustment of single-cell data, such as reclustering or refining annotations. This limitation hinders dynamic and accurate analysis of single-cell data, which often requires numerous interactive adjustments for re-labeling cell annotations.

To address this computational bottleneck, BioTuring developed AlphaSC, a comprehensive suite of fast and accurate algorithms to process single-cell data, leveraging the massive parallel power of GPU technology. In this report, we evaluated AlphaSC’s performance and accuracy against Seurat [2], Scanpy [3], and RAPIDS [1]. Our findings show that AlphaSC is significantly faster than both Seurat and Scanpy, achieving speeds more than a thousand times greater. Specifically, AlphaSC completed processing a 1.7 million-cell dataset in just 27 seconds, while Seurat required 29 hours for the same task.

Compared to RAPIDS [1], NVIDIA’s GPU-utilizing pipeline, AlphaSC not only demonstrates superior speed, being ten times faster, but also significantly reduces memory usage, both RAM and GPU memory. Importantly, AlphaSC maintains high accuracy consistently in all benchmarked scenarios, despite its rapid processing capabilities.

## 2 Results

We benchmark AlphaSC against Scanpy [3], Seurat [2], and RAPIDS [1] on 12 scRNAseq datasets with sizes ranging from 2,000 cells to 1.7 million cells on 5 steps: PCA, k-NN, t-SNE, UMAP and Louvain clustering.

### 2.1 Dataset description

We use 12 scRNAseq datasets, including Pancreas_2K [4], MDSC_4K [5], Glioblastoma_22K [6], Tcell_43K [7], Breast-Cancer_50K [8], Airways_77K [9], Epidermis_92K [10], Adrenal_387K [11], TabulaSapiens_481K [12], Lung_584K [13], Cerebellum_1M [11], and Cerebrum_1.7M [11].

Table 1 describes the details of the benchmark datasets, including the datasets’ IDs, number of cells, number of genes, and number of non-zero elements in the cell-by-gene matrix.

**Table 1:** Datasets used for benchmarking AlphaSC against Scanpy, Seurat, and RAPIDS

| Study ID | Number of Cells | Number of Genes | Number of Non-zero Elements |
| --- | --- | --- | --- |
| Pancreas_2K | 2,209 | 22,755 | 13,749,308 |
| MDSC_4K | 4,705 | 14,565 | 2,009,584 |
| Glioblastoma_22K | 22,637 | 17,128 | 46,429,854 |
| Tcell_43K | 43,112 | 19,993 | 143,930,315 |
| BreastCancer_50K | 50,693 | 21,962 | 74,598,494 |
| Airways_77K | 77,969 | 26,266 | 146,609,751 |
| Epidermis_92K | 92,889 | 2,468 | 229,242,565 |
| Adrenal_387K | 387,771 | 39,664 | 186,803,648 |
| TabulaSapiens_481K | 481,120 | 48,505 | 1,214,460,964 |
| Lung_584K | 584,884 | 26,187 | 1,140,188,601 |
| Cerebellum_1M | 1,092,000 | 44,330 | 782,448,367 |
| Cerebrum_1.7M | 1,751,246 | 45,526 | 1,237,849,906 |

### 2.2 Benchmark environment

The benchmark was performed on an NVIDIA DGX system with 8 NVIDIA A100 80 GB GPUs, 2 TB of RAM, and 256 CPU cores (2.45 GHz). The system operated on Ubuntu 22.04 LTS, with CUDA version 11.8. The high configuration of the server serves the purpose of running Scanpy, Seurat, and RAPIDS, while AlphaSC can run on low-cost servers with a 12 GB memory GPU.

We used Python version 3.10.12 and R version 4.4.0. We installed Scanpy version 1.9.5, AlphaSC version 0.6.0, and RAPIDS version 23.06 on the Python environment. On the R environment, we installed Seurat version 4.3.0.

### 2.3 End-to-end data processing pipeline benchmark

We benchmarked AlphaSC against Scanpy, Seurat, and RAPIDS. We use these four packages to perform an end-to-end scRNAseq data processing pipeline, including PCA, k-NN, t-SNE, UMAP, and Louvain clustering. Here, we present the total runtime and the computational resources cost of running the pipeline for each package. Detailed settings, runtime, computational resources cost, and results comparison for each analysis in the pipeline are described in the next sections.

The benchmark results are shown in Table 2. AlphaSC consistently shows its significant advantages in speed, RAM, and GPU memory consumption. On a dataset of size 1.7 million cells, AlphaSC finishes in 27 seconds, while RAPIDS takes 320 seconds, Seurat takes 106,886 seconds, and Scanpy takes 39,611 seconds. Notably, AlphaSC just consumes 3.2 GB GPU memory, and 17.2 RAM, while RAPIDS consumes 26.7 GB GPU and 41.1 GB memory. On the same dataset, Scanpy uses 38.8 GB and Seurat uses 137.4 GB RAM. Among these 4 tools, AlphaSC and RAPIDS use GPU, and their comparison is also depicted in Fig. 1.

**Table 2:** Comparative analysis of Scanpy, Seurat, AlphaSC, and RAPIDS on 12 scRNAseq datasets, showing total runtime (in seconds), maximum RAM and GPU memory consumption in gigabytes (denoted as RAM | GPU). Values are rounded to one decimal place.

| Study ID | Scanpy |  | Seurat |  | AlphaSC |  | RAPIDS |  |
| --- | --- | --- | --- | --- | --- | --- | --- | --- |
|  | Runtime | RAM | Runtime | RAM | Runtime | RAM\|GPU | Runtime | RAM\|GPU |
| Pancreas_2K | 30.6 | **0.4** | 26.1 | 1.4 | **1.9** | 0.9 \| **0.0** | 7.4 | 2.9 \| 1.3 |
| MDSC_4K | 40.7 | **0.5** | 58.3 | 1.1 | **1.8** | 0.8 \| **0.0** | 7.6 | 2.9 \| 1.4 |
| Glioblastoma_22K | 170.4 | **1.0** | 155.1 | 3.4 | **2.9** | 1.2 \| **0.1** | 8.6 | 3.0 \| 1.4 |
| Tcell_43K | 283.6 | **2.4** | 314.8 | 7.8 | **2.5** | 2.5 \| **0.2** | 9.8 | 4.1 \| 1.4 |
| BreastCancer_50K | 351.8 | 1.7 | 333.1 | 5.0 | **2.4** | **1.5** \| **0.2** | 9.5 | 3.7 \| 1.4 |
| Airways_77K | 522.4 | 2.7 | 620.3 | 8.1 | **3.7** | **2.6** \| **0.3** | 12.3 | 4.9 \| 2.1 |
| Epidermis_92K | 561.0 | **4.7** | 587.6 | 15.9 | **5.1** | 7.1 \| **1.4** | 11.6 | 5.7 \| 1.5 |
| Adrenal_387K | 4640.5 | 9.9 | 10502.6 | 24.0 | **7.0** | **3.2** \| **1.1** | 38.4 | 10.6 \| 6.3 |
| TabulaSapiens_481K | 4866.2 | 17.4 | 19842.5 | 59.6 | **9.2** | **16.9** \| **1.0** | 42.8 | 19.0 \| 5.1 |
| Lung_584K | 6774.1 | 18.6 | 7924.0 | 59.7 | **9.7** | **16.7** \| **1.7** | 57.9 | 23.3 \| 5.9 |
| Cerebellum_1M | 15425.0 | 24.4 | 42194.3 | 71.4 | **18.2** | **12.5** \| **2.5** | 156.2 | 32.5 \| 9.7 |
| Cerebrum_1.7M | 39610.7 | 38.8 | 106885.9 | 137.4 | **27.2** | **17.2** \| **3.2** | 320.4 | 41.1 \| 26.7 |

**Figure 1:**
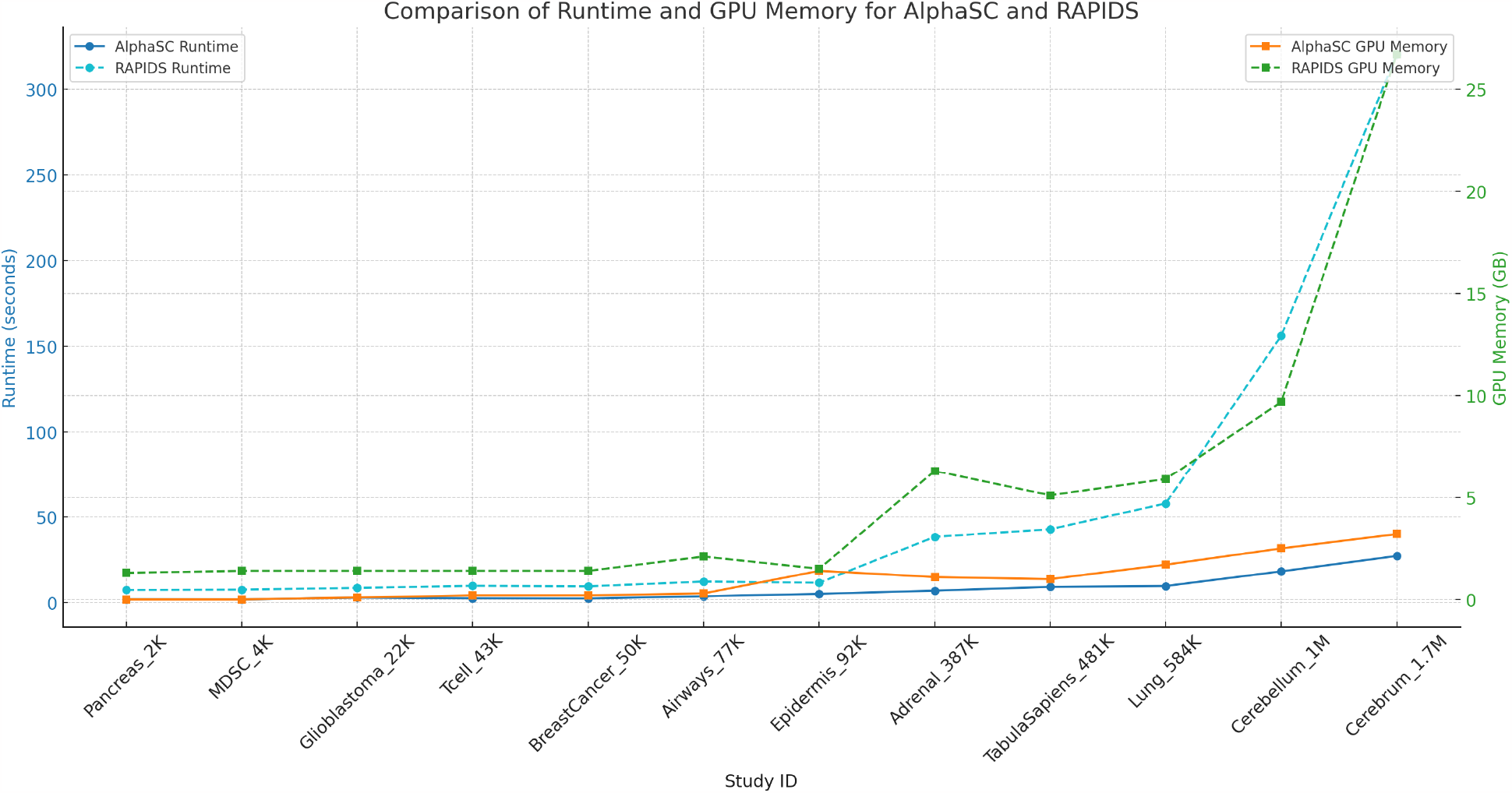
Comparative Analysis of AlphaSC and RAPIDS Across 12 scRNAseq Datasets. This graph presents a dual-axis comparison, illustrating both the runtime (seconds) and GPU memory usage (gigabytes) for each tool. While AlphaSC is an order of magnitude faster than RAPIDS, it also consumes much less GPU memory.

**Figure 2:**
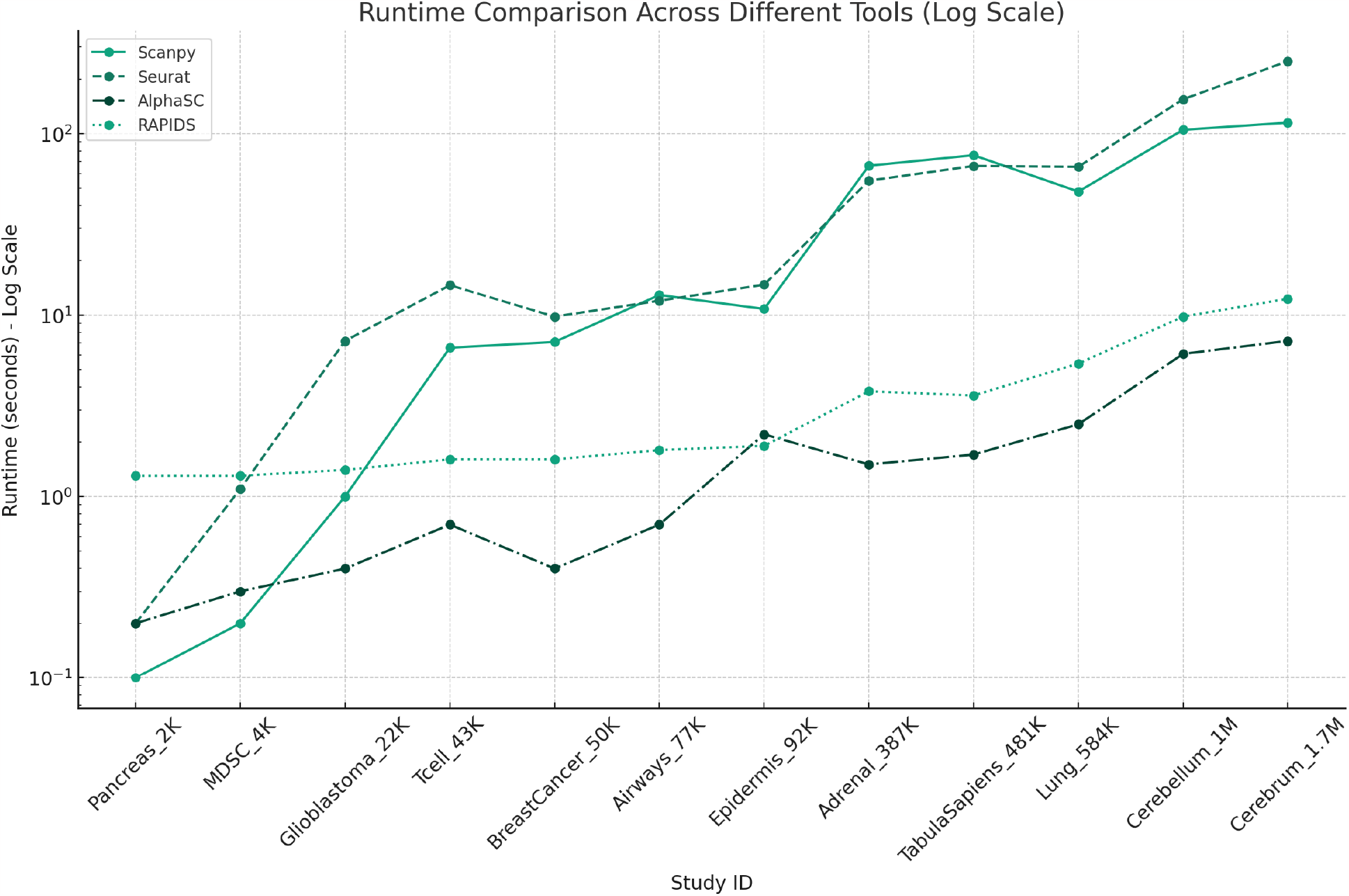
PCA Runtime. This plot illustrates the runtime performance of Scanpy, Seurat, AlphaSC, and RAPIDS across a range of single-cell RNA sequencing datasets, varying from 2K to 1.7M cells. The y-axis, displaying runtime in seconds, is set to a logarithmic scale to accommodate the wide span of runtime values.

### 2.4 PCA

We first use the function `FindVariableFeatures` from Seurat on the raw count matrix of each dataset to find the highly variable genes. We set `nfeatures=2000`, and other parameters are kept as default. Then, we filter the raw count matrix by selecting the top 2000 highly variable genes only. Finally, we scale the count matrix to obtain a z-score and apply a cutoff value of 10 standard deviations using the function `ScaleData` from Seurat with default parameters.

After filtering the matrix by the highly variable genes and scaling the filtered matrix, we perform PCA using Scanpy, Seurat, RAPIDS, and AlphaSC with the following functions and settings:

- Scanpy: `scanpy.tl.pca(n_comps=50)`. Other parameters are kept as default.
- Seurat: `RunPCA(npcs=50)`. Other parameters are kept as default.
- RAPIDS: `cuml.decomposition.PCA(n_components=50)`. Other parameters are kept as default.
- AlphaSC: `bioalpha.singlecell.preprocessing.pca(n_comps=50)`. Other parameters are kept as default.

The benchmark results are shown in Table 3. On average, AlphaSC runs 18 times faster than Scanpy, 27 times faster than Seurat, and 2 times faster than RAPIDS. The total variance explained produced by all packages are highly similar, and all are over >99% similar to the results obtained using Seurat.

**Table 3:** Comprehensive analysis of PCA runtime in seconds and RAM consumption (GB) for Scanpy, Seurat, AlphaSC, and RAPIDS across 12 scRNAseq datasets. For RAPIDS and AlphaSC, GPU memory (GB) is also shown.

| Study ID | Scanpy |  | Seurat |  | AlphaSC |  | RAPIDS |  |
| --- | --- | --- | --- | --- | --- | --- | --- | --- |
|  | *Runtime* | *RAM* | *Runtime* | *RAM* | *Runtime* | *RAM\|GPU* | *Runtime* | *RAM\|GPU* |
| Pancreas_2K | **0.1** | **0.4** | 0.2 | 1.4 | 0.2 | 0.9 \| **0.0** | 1.3 | 2.4 \| 0.6 |
| MDSC_4K | **0.2** | **0.3** | 1.1 | 1.1 | 0.3 | 0.8 \| **0.0** | 1.3 | 2.3 \| 0.6 |
| Glioblastoma_22K | 1.0 | **1.0** | 7.2 | 3.4 | **0.4** | 1.2 \| **0.1** | 1.4 | 3.0 \| 0.9 |
| Tcell_43K | 6.6 | **2.4** | 14.6 | 7.8 | **0.7** | 2.5 \| **0.2** | 1.6 | 4.1 \| 1.2 |
| BreastCancer_50K | 7.1 | 1.7 | 9.8 | 5.0 | **0.4** | **1.5 \| 0.1** | 1.6 | 3.7 \| 1.3 |
| Airways_77K | 12.9 | 2.7 | 12.0 | 8.1 | **0.7** | **2.6 \| 0.1** | 1.8 | 4.9 \| 2.1 |
| Epidermis_92K | 10.8 | **4.7** | 14.7 | 15.9 | 2.2 | 7.1 \| **1.4** | **1.9** | 5.7 \| 1.2 |
| Adrenal_387K | 66.5 | 8.2 | 55.0 | 24.0 | **1.5** | **3.2 \| 0.1** | 3.8 | 10.6 \| 6.3 |
| TabulaSapiens_481K | 76.0 | 17.4 | 66.3 | 59.6 | **1.7** | **16.9 \| 0.1** | 3.6 | 19.0 \| 5.1 |
| Lung_584K | 47.9 | 18.6 | 65.6 | 59.7 | **2.5** | **16.7 \| 0.6** | 5.4 | 23.3 \| 5.9 |
| Cerebellum_1M | 104.8 | 24.4 | 154.5 | 71.4 | **6.1** | **12.5 \| 0.6** | 9.8 | 32.5 \| 9.7 |
| Cerebrum_1.7M | 114.8 | 38.8 | 250.6 | 137.4 | **7.2** | **17.2 \| 0.9** | 12.3 | 41.1 \| 26.7 |

### 2.5 k-NN

We first use the function `FindVariableFeatures` from Seurat on the raw count matrix of each dataset to find the highly variable genes. We set `nfeatures=2000`, and other parameters are kept as default. Then, we filter the raw count matrix by selecting the top 2000 highly variable genes only. Finally, we scale the count matrix to obtain a z-score and apply a cutoff value of 10 standard deviations using the function `ScaleData` from Seurat with default parameters.

After filtering the matrix by the highly variable genes and scaling the filtered matrix, we perform PCA using the function `RunPCA` from Seurat with `npcs=50`. Other parameters are kept as default. Then, we use the PCA matrix as input to perform k-NN using Scanpy, Seurat, RAPIDS, and AlphaSC with the following functions and settings:

- Scanpy: `scanpy.pp.neighbors(k=30)`. Other parameters are kept as default.
- Seurat: `FindNeighbors(k=30)`. Other parameters are kept as default.
- RAPIDS: `cuml.neighbors.NearestNeighbors(n_neighbors=30)`. Other parameters are kept as default.
- AlphaSC: `bioalpha.singlecell.preprocessing.neighbors(k=30)`. Other parameters are kept as default.

The k-NN performance benchmark results are shown in Table 4, and Figure 3. On average, AlphaSC runs 195 times faster than Scanpy, 247 times faster than Seurat, and 11 times faster than RAPIDS.

**Table 4:** Comprehensive analysis of k-NN performance (runtime and RAM) for Scanpy, Seurat, AlphaSC, and RAPIDS across 12 scRNAseq datasets. For RAPIDS and AlphaSC, GPU memory consumption (GB) is also included.

| Study ID | Scanpy |  | Seurat |  | AlphaSC |  | RAPIDS |  |
| --- | --- | --- | --- | --- | --- | --- | --- | --- |
|  | *Runtime* | *RAM* | *Runtime* | *RAM* | *Runtime* | *RAM\|GPU* | *Runtime* | *RAM\|GPU* |
| Pancreas_2K | 12.4 | **0.4** | 0.6 | 0.4 | **0.3** | 0.5 \| **0.0** | 1.2 | 2.2 \| 1.1 |
| MDSC_4K | 7.6 | 0.5 | 1.2 | **0.4** | **0.3** | 0.5 \| **0.0** | 1.2 | 2.2 \| 1.0 |
| Glioblastoma_22K | 36.4 | 0.7 | 6.3 | **0.5** | **0.3** | 0.6 \| **0.1** | 1.2 | 2.2 \| 1.0 |
| Tcell_43K | 40.8 | 0.8 | 12.2 | 0.7 | **0.3** | **0.6 \| 0.1** | 1.3 | 2.3 \| 0.9 |
| BreastCancer_50K | 43.6 | 0.9 | 14.8 | 0.7 | **0.4** | **0.6 \| 0.1** | 1.3 | 2.3 \| 1.0 |
| Airways_77K | 51.1 | 1.1 | 25.7 | 0.8 | **0.4** | **0.7 \| 0.1** | 1.5 | 2.3 \| 1.2 |
| Epidermis_92K | 54.8 | 1.2 | 28.4 | 0.8 | **0.3** | **0.7 \| 0.2** | 1.4 | 2.3 \| 1.0 |
| Adrenal_387K | 148.6 | 3.3 | 172.3 | 1.9 | **0.8** | **1.5 \| 0.6** | 4.0 | 2.8 \| 1.4 |
| TabulaSapiens_481K | 175.4 | 4.0 | 197.9 | 2.3 | **1.0** | **1.8 \| 0.8** | 5.5 | 2.8 \| 1.3 |
| Lung_584K | 209.2 | 4.7 | 254.6 | 2.6 | **0.9** | **2.0 \| 0.9** | 7.4 | 2.9 \| 1.4 |
| Cerebellum_1M | 357.7 | 9.9 | 534.7 | 4.5 | **1.5** | **3.4 \| 1.7** | 22.8 | 3.5 \| **1.5** |
| Cerebrum_1.7M | 582.6 | 16.4 | 931.7 | 7.1 | **2.3** | 5.2 \| 2.7 | 55.6 | **4.5 \| 2.4** |

**Figure 3:**
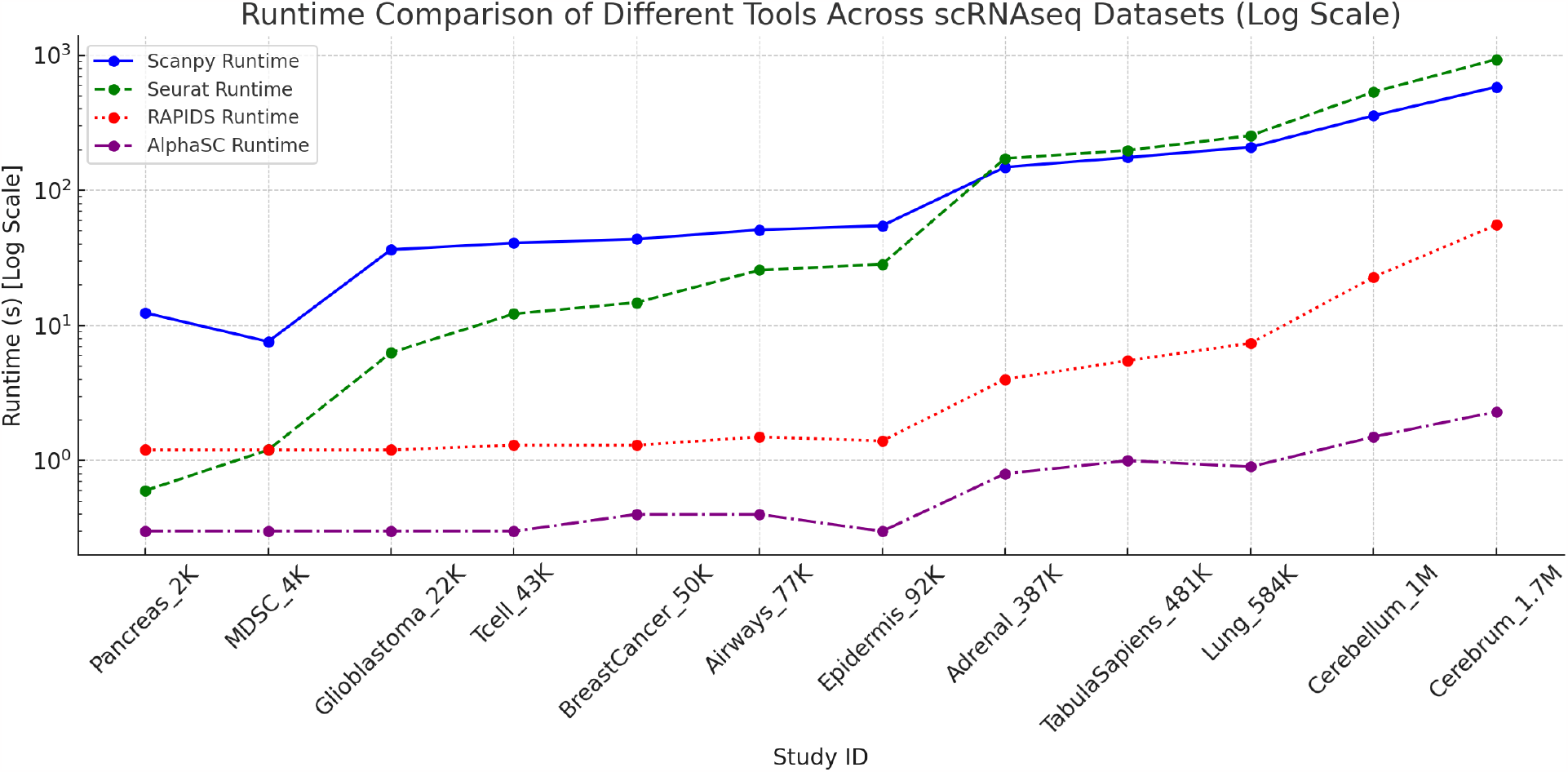
k-NN Performance. This plot illustrates the runtime performance of Scanpy, Seurat, AlphaSC, and RAPIDS across a range of single-cell RNA sequencing datasets, varying from 2K to 1.7M cells. The y-axis, displaying runtime in seconds, is set to a logarithmic scale to accommodate the wide span of runtime values.

We evaluate the resulting k-NN graphs by first constructing the exact k-NN graphs using KDTree [14] and calculating the accuracy score between the exact k-NN graphs and the k-NN graphs produced by the four packages. The pseudocode to calculate the accuracy score between an exact k-NN graph and an approximate k-NN graph is described in Algorithm 1.

Table 5 shows the accuracy scores of the 4 tools. Seurat has low accuracy scores on several datasets. On the other hand, AlphaSC achieves the optimal accuracy scores on all benchmarked datasets.

**Table 5:** Average accuracy score between the exact k-NN graph and the corresponding k-NN graph produced by Scanpy, Seurat, RAPIDS, and AlphaSC, respectively.

| Study ID | Scanpy | Seurat | RAPIDS | AlphaSC |
| --- | --- | --- | --- | --- |
| Pancreas_2K | 1.00 | 0.96 | 1.00 | 1.00 |
| MDSC_4K | 1.00 | 0.79 | 1.00 | 1.00 |
| Glioblastoma_22K | 1.00 | 0.95 | 1.00 | 1.00 |
| Tcell_43K | 1.00 | 0.89 | 1.00 | 1.00 |
| BreastCancer_50K | 1.00 | 0.94 | 1.00 | 1.00 |
| Airways_77K | 0.99 | 0.86 | 1.00 | 1.00 |
| Epidermis_92K | 1.00 | 0.99 | 1.00 | 1.00 |
| Adrenal_387K | 0.99 | 0.77 | 1.00 | 1.00 |
| TabulaSapiens_481K | 0.97 | 0.67 | 1.00 | 1.00 |
| Lung_584K | 1.00 | 0.93 | 1.00 | 1.00 |
| Cerebellum_1M | 0.99 | 0.80 | 1.00 | 1.00 |
| Cerebrum_1.7M | 0.98 | 0.72 | 1.00 | 1.00 |

#### Algorithm 1 Pseudocode for calculating the accuracy score between an exact k-NN graph and an approximate k-NN graph

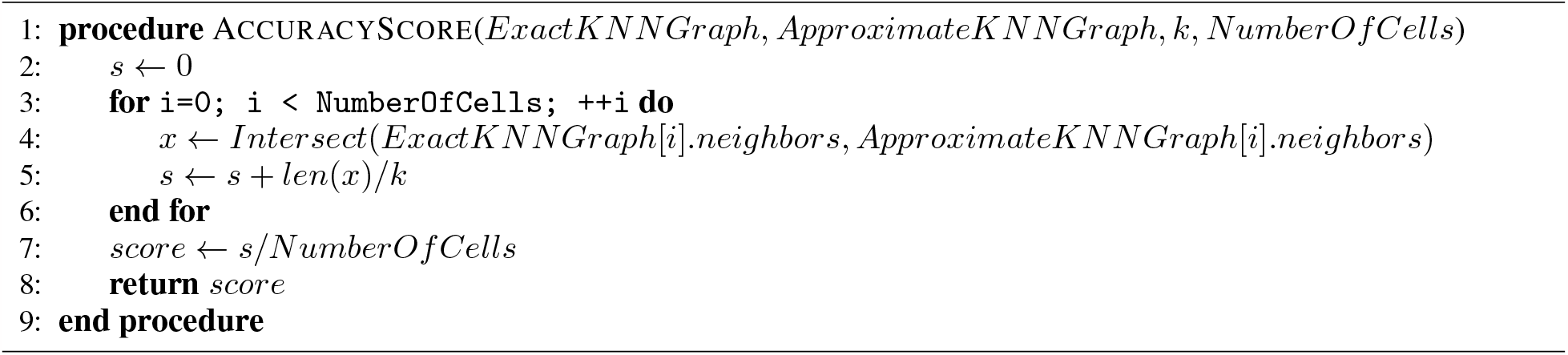

### 2.6 t-SNE

We first use the function `FindVariableFeatures` from Seurat on the raw count matrix of each dataset to find the highly variable genes. We set `nfeatures=2000`, and other parameters are kept as default. Then, we filter the raw count matrix by selecting the top 2000 highly variable genes only. Finally, we scale the count matrix to obtain a z-score and apply a cutoff value of 10 standard deviations using the function `ScaleData` from Seurat with default parameters.

After filtering the matrix by the highly variable genes and scaling the filtered matrix, we perform PCA using the function `RunPCA` from Seurat with `npcs=50`. Other parameters are kept as default. Then, we use the PCA matrix as input to perform t-SNE using scanpy, Seurat, RAPIDS, and AlphaSC with the following functions and settings:

- scanpy: `scanpy.tl.tsne(perplexity=30)`. Other parameters are kept as default.
- Seurat: `RunTSNE(perplexity=30)`. Other parameters are kept as default.
- RAPIDS: `cuml.TSNE(n_neighbors=30)`. Other parameters are kept as default.
- AlphaSC: `bioalpha.singlecell.tools.tsne(perplexity=30)`. Other parameters are kept as default.

The benchmark results are shown in Table 6, Figure 4. On average, AlphaSC runs 423 times faster than scanpy, 5306 times faster than Seurat, and 6 times faster than RAPIDS.

**Table 6:** Combined analysis of t-SNE runtime and RAM/GPU consumption for Scanpy, Seurat, AlphaSC, and RAPIDS on 12 scRNAseq datasets.

| Study ID | Scanpy |  | Seurat |  | AlphaSC |  | RAPIDS |  |
| --- | --- | --- | --- | --- | --- | --- | --- | --- |
|  | *Runtime* | *RAM* | *Runtime* | *RAM* | *Runtime* | *RAM\|GPU* | *Runtime* | *RAM\|GPU* |
| Pancreas_2K | 5.0 | **0.3** | 5.0 | 0.4 | **0.7** | 0.5 \| **0.0** | 1.6 | 2.2 \| 0.6 |
| MDSC_4K | 9.6 | **0.3** | 12.2 | 0.4 | **0.6** | 0.5 \| **0.0** | 1.7 | 2.2 \| 1.0 |
| Glioblastoma_22K | 39.6 | **0.4** | 74.1 | 0.5 | **1.5** | 0.6 \| **0.1** | 2.0 | 2.2 \| 1.0 |
| Tcell_43K | 86.1 | **0.5** | 184.6 | 0.7 | **0.7** | 0.6 \| **0.1** | 2.3 | 2.3 \| 1.0 |
| BreastCancer_50K | 90.8 | **0.6** | 187.0 | 0.7 | **0.8** | **0.6 \| 0.1** | 2.2 | 2.3 \| 1.0 |
| Airways_77K | 149.8 | 0.7 | 372.0 | 0.9 | **1.7** | **0.6 \| 0.1** | 2.6 | 2.3 \| 1.1 |
| Epidermis_92K | 158.9 | **0.7** | 322.0 | 0.9 | **1.6** | **0.7 \| 0.2** | 2.4 | 2.3 \| 1.0 |
| Adrenal_387K | 958.2 | 2.5 | 8528.8 | 2.5 | **2.2** | **1.1 \| 0.6** | 9.3 | 2.5 \| 1.2 |
| TabulaSapiens_481K | 1009.6 | 3.0 | 18199.6 | 3.0 | **3.6** | **1.2 \| 0.8** | 16.2 | 2.5 \| 1.2 |
| Lung_584K | 1536.9 | 3.2 | 4552.5 | 3.6 | **3.1** | **1.3 \| 0.9** | 14.8 | 2.5 \| 1.2 |
| Cerebellum_1M | 3032.5 | 6.3 | 34284.8 | 6.4 | **5.1** | **2.0 \| 1.7** | 42.4 | 2.8 \| **1.3** |
| Cerebrum_1.7M | 5930.1 | 9.9 | 96425.4 | 10.1 | **9.2** | **3.0 \| 2.7** | 96.6 | 3.2 \| **1.5** |

**Figure 4:**
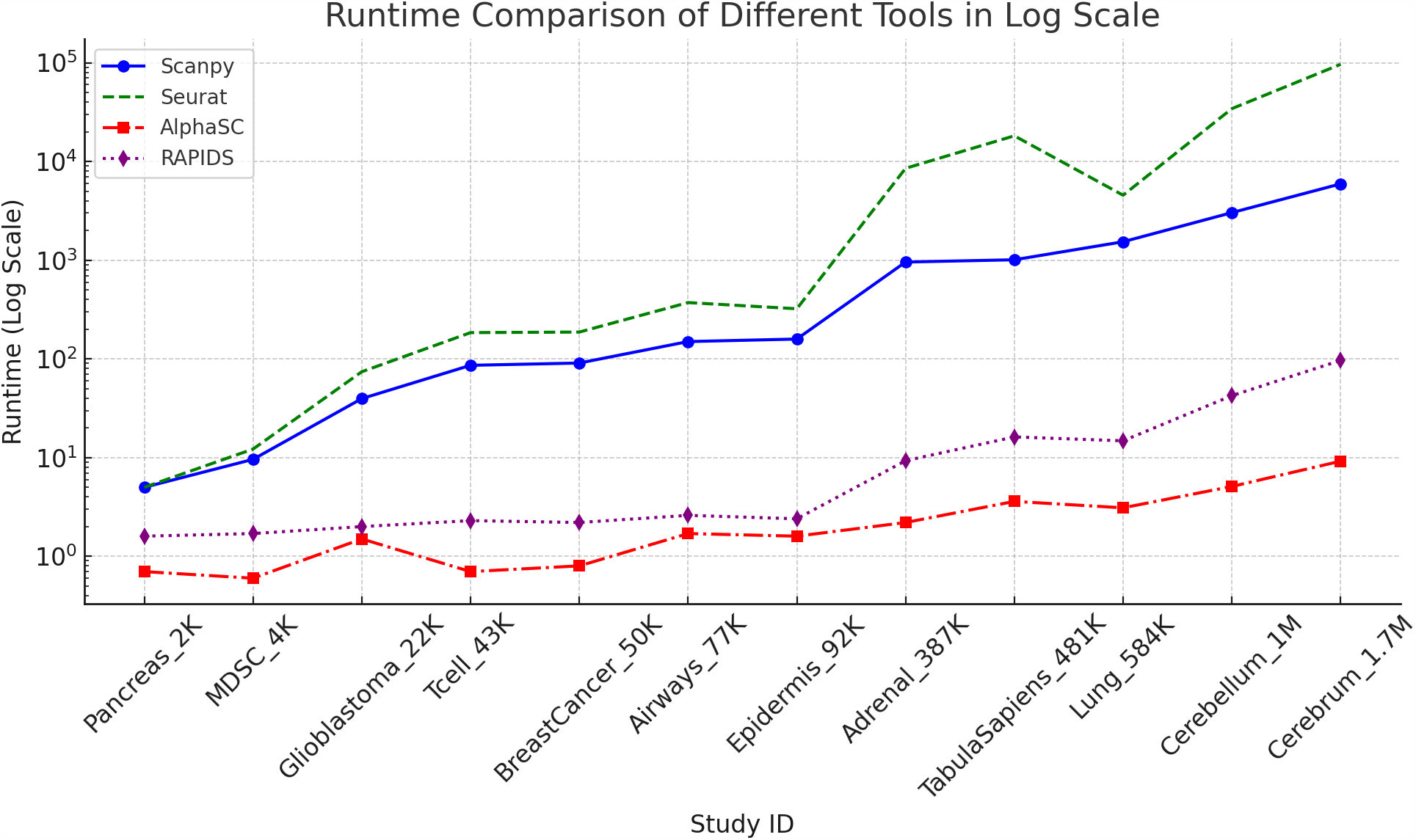
t-SNE Performance. This plot illustrates the t-SNE runtime performance of Scanpy, Seurat, AlphaSC, and RAPIDS across datasets. The y-axis, displaying runtime in seconds, is set to a logarithmic scale to accommodate the wide span of runtime values.

### 2.7 UMAP

We first use the function `FindVariableFeatures` from Seurat on the raw count matrix of each dataset to find the highly variable genes. We set `nfeatures=2000`, and other parameters are kept as default. Then, we filter the raw count matrix by selecting the top 2000 highly variable genes only. Finally, we scale the count matrix to obtain a z-score and apply a cutoff value of 10 standard deviations using the function `ScaleData` from Seurat with default parameters.

After filtering the matrix by the highly variable genes and scaling the filtered matrix, we perform PCA using the function `RunPCA` from Seurat with `npcs=50`. Other parameters are kept as default. Then, we use the PCA matrix as input to perform UMAP using scanpy, Seurat, RAPIDS, and AlphaSC with the following functions and settings:

- scanpy: `scanpy.tl.umap()`. All parameters are kept as default.
- Seurat: `RunUMAP()`. All parameters are kept as default.
- RAPIDS: `cuml.UMAP()`. All parameters are kept as default.
- AlphaSC: `bioalpha.singlecell.tools.umap()`. All parameters are kept as default.

The benchmark results are shown in Table 7, Figure 5. On average, AlphaSC runs 287 times faster than scanpy, 276 times faster than Seurat, and 6 times faster than RAPIDS.

**Table 7:** Combined analysis of UMAP runtime and RAM/GPU consumption for Scanpy, Seurat, AlphaSC, and RAPIDS on 12 scRNAseq datasets.

| Study ID | Scanpy |  | Seurat |  | AlphaSC |  | RAPIDS |  |
| --- | --- | --- | --- | --- | --- | --- | --- | --- |
|  | *Runtime* | *RAM* | *Runtime* | *RAM* | *Runtime* | *RAM\|GPU* | *Runtime* | *RAM\|GPU* |
| Pancreas_2K | 11.4 | **0.4** | 19.9 | 0.7 | **0.5** | 0.8 \| **0.0** | 1.4 | 2.2 \| 1.1 |
| MDSC_4K | 15.6 | **0.4** | 41.7 | 0.8 | **0.3** | 0.8 \| **0.0** | 1.4 | 2.2 \| 0.9 |
| Glioblastoma_22K | 59.1 | **0.8** | 52.8 | 1.1 | **0.4** | **0.8 \| 0.0** | 1.4 | 2.2 \| 1.0 |
| Tcell_43K | 74.4 | **0.9** | 74.8 | 1.2 | **0.5** | **0.9 \| 0.1** | 1.5 | 2.3 \| 1.0 |
| BreastCancer_50K | 77.8 | 1.0 | 77.7 | 1.4 | **0.5** | **0.9 \| 0.1** | 1.6 | 2.3 \| 1.2 |
| Airways_77K | 118.6 | 1.3 | 101.3 | 1.8 | **0.6** | **1.0 \| 0.1** | 1.7 | 2.3 \| 1.0 |
| Epidermis_92K | 113.7 | 1.4 | 105.6 | 1.9 | **0.6** | **1.0 \| 0.1** | 1.9 | 2.3 \| 1.0 |
| Adrenal_387K | 469.3 | 4.5 | 424.3 | 5.7 | **2.0** | **1.5 \| 0.5** | 5.6 | 2.5 \| 1.3 |
| TabulaSapiens_481K | 565.9 | 5.6 | 532.8 | 7.2 | **2.4** | **1.7 \| 0.6** | 7.6 | 2.5 \| 1.2 |
| Lung_584K | 679.8 | 7.6 | 625.2 | 8.8 | **2.5** | **1.8 \| 0.7** | 10.5 | 2.5 \| 1.4 |
| Cerebellum_1M | 1460.6 | 15.8 | 1649.5 | 19.2 | **4.4** | 3.2 \| **1.3** | 29.6 | **2.8 \| 1.3** |
| Cerebrum_1.7M | 2617.5 | 27.6 | 2322.5 | 35.2 | **7.1** | **4.9 \| 2.2** | 71.1 | 3.2 \| **2.0** |

**Figure 5:**
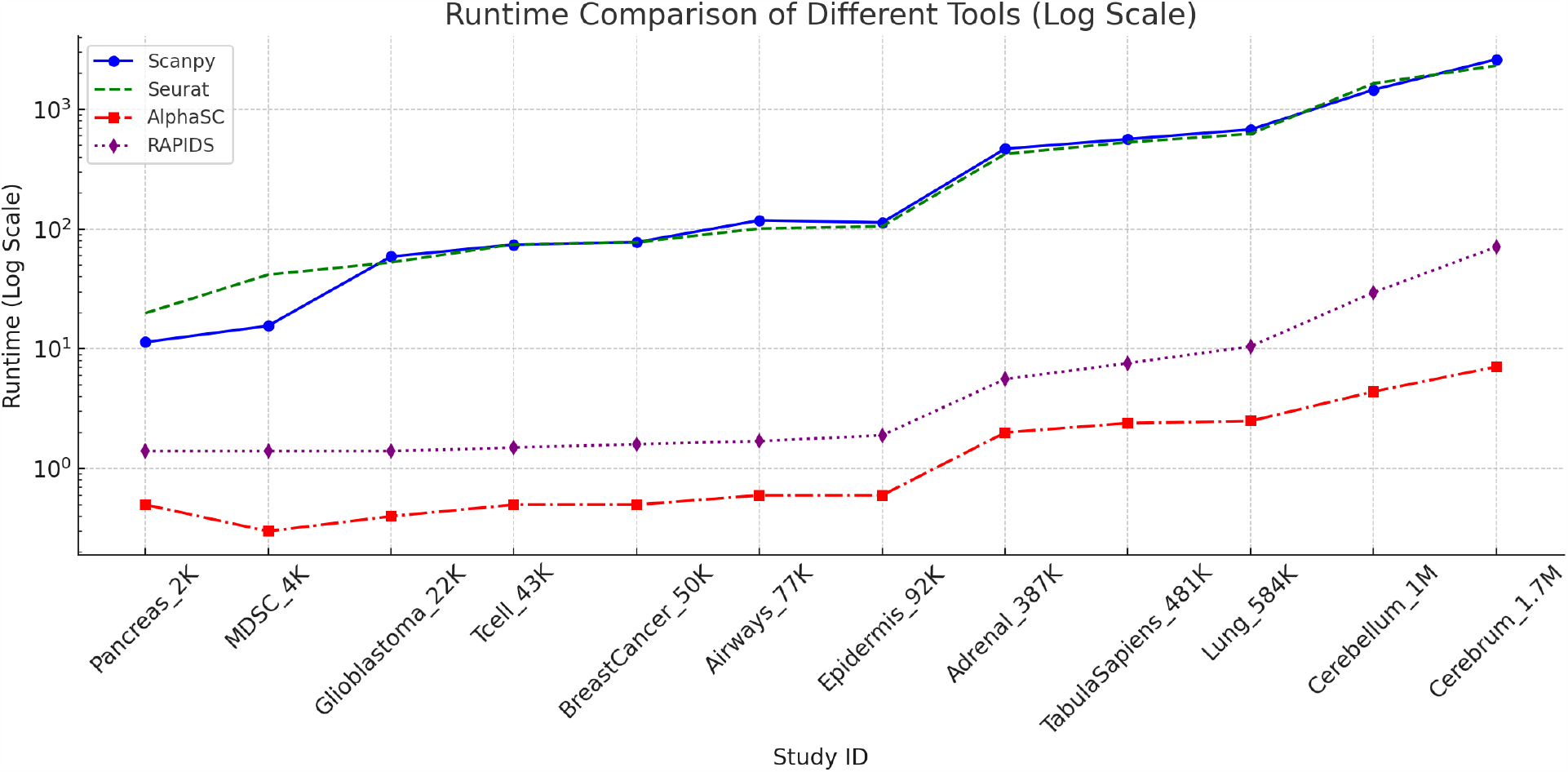
UMAP Performance. This plot illustrates the runtime performance of UMAP implementation on Scanpy, Seurat, AlphaSC, and RAPIDS across a range of single-cell RNA sequencing datasets, varying from 2K to 1.7M cells. The y-axis, displaying runtime in seconds, is set to a logarithmic scale to accommodate the wide span of runtime values.

### 2.8 Louvain clustering

We first use the function `FindVariableFeatures` from Seurat on the raw count matrix of each dataset to find the highly variable genes. We set `nfeatures=2000`, and other parameters are kept as default. Then, we filter the raw count matrix by selecting the top 2000 highly variable genes only. Finally, we scale the count matrix to obtain a z-score and apply a cutoff value of 10 standard deviations using the function `ScaleData` from Seurat with default parameters.

After filtering the matrix by the highly variable genes and scaling the filtered matrix, we perform PCA using the function `RunPCA` from Seurat with `npcs=50`. Other parameters are kept as default. Then, we use the PCA matrix as input to perform Louvain clustering using scanpy, Seurat, RAPIDS, and AlphaSC with the following functions and settings:

- Scanpy: `scanpy.tl.louvain(resolution=1)`. Other parameters are kept as default.
- Seurat: `FindClusters(resolution=1)`. Other parameters are kept as default.
- RAPIDS: `cugraph.louvain(resolution=1)`. Other parameters are kept as default.
- AlphaSC: `bioalpha.singlecell.tools.louvain(resolution=1)`. Other parameters are kept as default.

The benchmark results are shown in Table 8, and Figure 6. On average, AlphaSC runs 8175 times faster than scanpy, 2749 times faster than Seurat, and 32 times faster than RAPIDS.

**Table 8:**
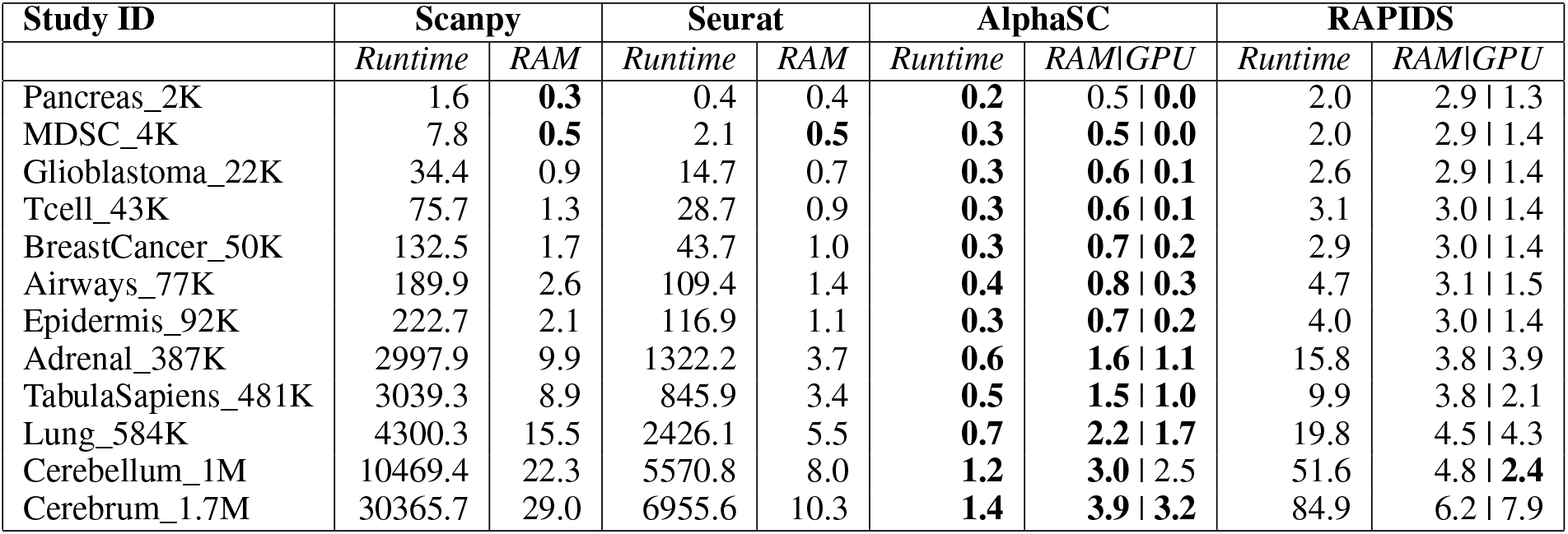
Combined analysis of Louvain clustering runtime and RAM/GPU consumption for Scanpy, Seurat, AlphaSC, and RAPIDS on 12 scRNAseq datasets.

**Figure 6:**
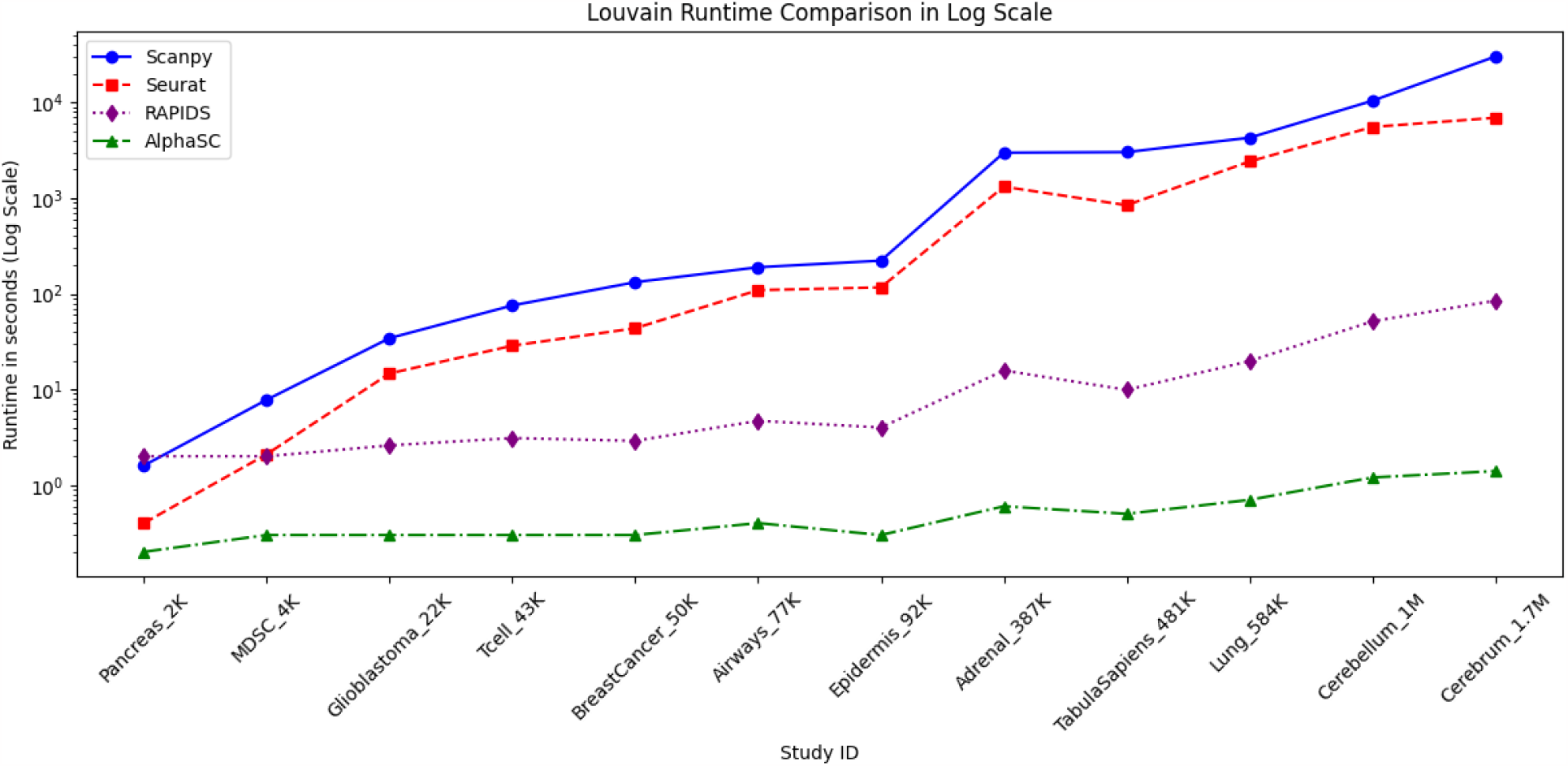
Louvain Performance. This plot illustrates the runtime performance of Louvain implementation on Scanpy, Seurat, RAPIDS, and AlphaSC across a range of single-cell RNA sequencing datasets, varying from 2K to 1.7M cells. The y-axis, displaying runtime in seconds, is set to a logarithmic scale to accommodate the wide span of runtime values.

To evaluate the quality of the clustering implementation in these tools, we use modularity scores [15]. Modularity score is a scalar value that measures the strength of division of a network into modules (also called communities). For a given division of the network, the modularity score compares the density of edges inside communities to the density of edges between communities. Higher modularity values indicate a stronger division of the network into communities. The modularity *Q* can be expressed as:

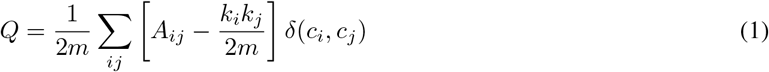

where *A*_*ij*_ represents the edge weight between nodes *i* and *j, k*_*i*_ and *k*_*j*_ are the sum of the weights of the edges attached to nodes *i* and *j, m* is the sum of all of the edge weights in the network, and *δ*(*c*_*i*_, *c*_*j*_) is the Kronecker delta function, which is 1 if *i* and *j* are in the same community and 0 otherwise.

The modularity scores of the four tools are provided in Table 9.

**Table 9:** Modularity scores calculated on Louvain clustering results of Scanpy, Seurat, AlphaSC, and RAPIDS on 12 scRNAseq datasets. The higher the score, the better.

| Study ID | scanpy | Seurat | AlphaSC | RAPIDS |
| --- | --- | --- | --- | --- |
| Pancreas_2K | 0.70 | 0.71 | **0.83** | 0.82 |
| MDSC_4K | **0.83** | **0.83** | 0.70 | 0.69 |
| Glioblastoma_22K | **0.87** | **0.87** | **0.87** | 0.86 |
| Tcell_43K | **0.90** | **0.90** | 0.83 | 0.83 |
| BreastCancer_50K | 0.84 | 0.84 | **0.90** | **0.90** |
| Airways_77K | **0.88** | **0.88** | **0.88** | **0.88** |
| Epidermis_92K | **0.92** | **0.92** | 0.90 | 0.90 |
| Adrenal_387K | 0.91 | 0.91 | **0.92** | 0.92 |
| TabulaSapiens_481K | 0.91 | 0.92 | **0.94** | 0.94 |
| Lung_584K | **0.94** | **0.94** | **0.94** | **0.94** |
| Cerebellum_1M | **0.94** | **0.94** | 0.91 | 0.91 |
| Cerebrum_1.7M | **0.92** | **0.92** | 0.91 | 0.91 |

## 3 Discussion

Through several benchmarks, we demonstrate AlphaSC is a powerful and efficient GPU-accelerated toolkit, providing significant performance and memory usage advantages over commonly used packages such as Scanpy, Seurat, and RAPIDS while being highly accurate. AlphaSC resolves the bottleneck for processing and analyzing multi-million cell datasets, enables researchers to process large-scale single-cell datasets interactively.

## Footnotes

1 AlphaSC enables real-time analysis of large single-cell datasets, and facilitates the creation of million-cell in-silico atlases. It also paves the way for generative AI models to efficiently access and analyze single-cell data from the BioTuring database.

